# Molecular identification of bats from Wavulgalge cave, Wellawaya, Sri Lanka

**DOI:** 10.1101/2021.12.19.473364

**Authors:** Thejanee Perera, Franziska Schwarz, Therese Muzeniek, Sahan Siriwardana, Beate Becker-Ziaja, Inoka C. Perera, Shiroma Handunnetti, Jagathpriya Weerasena, Gayani Premawansa, Sunil Premawansa, Andreas Nitsche, Wipula Yapa, Claudia Kohl

## Abstract

This is the first report on the molecular identification and phylogeny of *Rousettus leschenaultii, Rhinolophus rouxii, Hipposideros speoris, Hipposideros lankadiva, Miniopterus fuliginosus* bat species in Sri Lanka, inferred from *mitochondrially encoded cytochrome b* gene sequences. Wellawaya Wavulgalge cave in Sri Lanka is one of the largest sympatric colonies found on the island, occupied by five species of bats. Recent research has indicated that bats show enormous cryptic genetic diversity. Moreover, even in the same species, acoustic properties of echolocation calls and morphological features such as fur colour could vary in different populations. Therefore, we have used molecular techniques for the accurate identification of five bat species recorded in one of the largest cave populations in Sri Lanka. Bats were caught using a hand net and saliva samples were collected non-invasively from each bat using a sterile oral swab. Nucleic acids were extracted from oral swab samples and mitochondrial DNA was amplified using primers targeting the *mitochondrially encoded cytochrome b* gene. This study identified the bat species recorded in the Wellawaya cave as *Rousettus leschenaultii, Rhinolophus rouxii, Hipposideros speoris, Hipposideros lankadiva* and *Miniopterus fuliginosus*. Our findings will contribute to future conservation and systematic studies of bats in Sri Lanka. This study will also provide the basis for a genetic database of Sri Lankan bats.

## 1. Introduction

Bats (Order: Chiroptera) are the second largest group of mammals documented with 1,293 species in the world (1). Being a tropical country, with long day lengths and seasonal rainfall, Sri Lanka is endowed with high biological diversity. So far 30 species of bats have been recorded in Sri Lanka (2). Consequently, bats are the largest group of mammals recorded in Sri Lanka, accounting for approximately 1/3 of all mammalian species present on the island.

The order Chiroptera was traditionally subdivided into the two suborders, Megachiroptera and Microchiroptera, based on the evolution of echolocation (3). The Suborder Megachiroptera comprises frugivorous, predominantly the non-echolocating bats in a single-family *Pteropodidae*, whereas the most ecologically diverse of the two suborders, Microchiroptera consists of 16 families which includes echolocating bats with diverse feeding habits. However, more recently, the suborders Megachiroptera and Microchiroptera have been changed into two new suborders based on molecular phylogeny: Yinpterochiroptera and Yangochiroptera. All the Megachiropterans and the members of the superfamily Rhinolopoidea are grouped in the Suborder Yinpterochiroptera and the remaining microchiropteran species were included in the suborder Yangochiroptera (4). Housekeeping genes have low evolutionary rates and are therefore suitable for accurate species identification (5). These genes show low intraspecific variation and high interspecific variation. The *mitochondrially encoded cytochrome b* (MT-CYB) gene is a commonly used housekeeping gene that involves oxidative phosphorylation in cellular respiration. The whole gene sequence of the MT-CYB gene can be found in the National Center for Biotechnology Information Support Center (NCBI) database for most organisms.

Although some studies have been carried out on ecological and behavioural aspects of Sri Lankan bats, research on molecular identification is limited to a single species (6). Several bat species form a unique sympatric colony in the Wavulgalge cave in Wellawaya, Sri Lanka (7). Approximately 100,000 individual bats are inhabiting the cave. Based on external morphometric characteristics, these five species were identified as *Rousettus leschenaultii, Rhinolophus rouxii, Hipposideros speoris, Hipposideros lankadiva* and *Miniopterus fuliginosus*. In this study, amplification of the MT-CYB gene was utilized for the accurate identification of these five bat species inhabiting the Wavulgalge cave.

## 2. Materials and Methods

### Study

Investigative research on Sri Lankan bats was approved by the Department of Wildlife Conservation, Sri Lanka (permit No. WL/3/2/05/18) and conducted in accordance with the Fauna & Flora Protection Ordinance (FFPO) of Sri Lanka. Ethical clearance for the study was obtained from the Institute of Biology, Sri Lanka (ERC IOBSL 170 01 18). Samples were collected in a non-invasive manner. Animal capturing, handling and sampling was done in accordance with the guidelines of the American Society of Mammalogists for the use of wild mammals in research and education (8). The sampling site is a natural cave situated in Nikapitiya, Wellawaya (6°43’00.0”N 81°03’00.0”E) in Monaragala district, Sri Lanka. The roost is known as Wavulgalge cave. It is a lithophilic type of roost and one of the largest sympatric bat roosts found on the island. This underground cave expands over 45,000 m^2^ of area. The Wavulgalge cave provides a stable microclimate and protection from predators and extreme weather conditions (7). Sample collection was carried out during March and July of 2018 and in January 2019.

Adequate personal protective equipment such as safety gloves, safety glasses and FFP3 masks were used during animal capturing, handling and sample collection to reduce the potential risk of zoonotic or anthropozoonotic pathogen transmission.

Bats were captured using hand nets while they were emerging from the roost in the evenings. Captured live bats were placed into cotton bags and were kept in a cool dry place until further processing. Bat species were macroscopically identified using external morphological features (Fig 1.). Morphometric parameters and locational data were recorded in the data sheets (Fig 2.). Bats were released immediately after recording the morphometric data and the collection of samples, at the cave entrance. Saliva samples were collected from each bat using a sterile oral swab and snap-frozen using liquid nitrogen. Collected samples were transported to the laboratory in a dry-shipper and were stored at a −80°C freezer until further analysis. Laboratory analyses of the samples were carried out under BSL-2 conditions at the Robert Koch Institute, Berlin, Germany.

**Fig 1.**
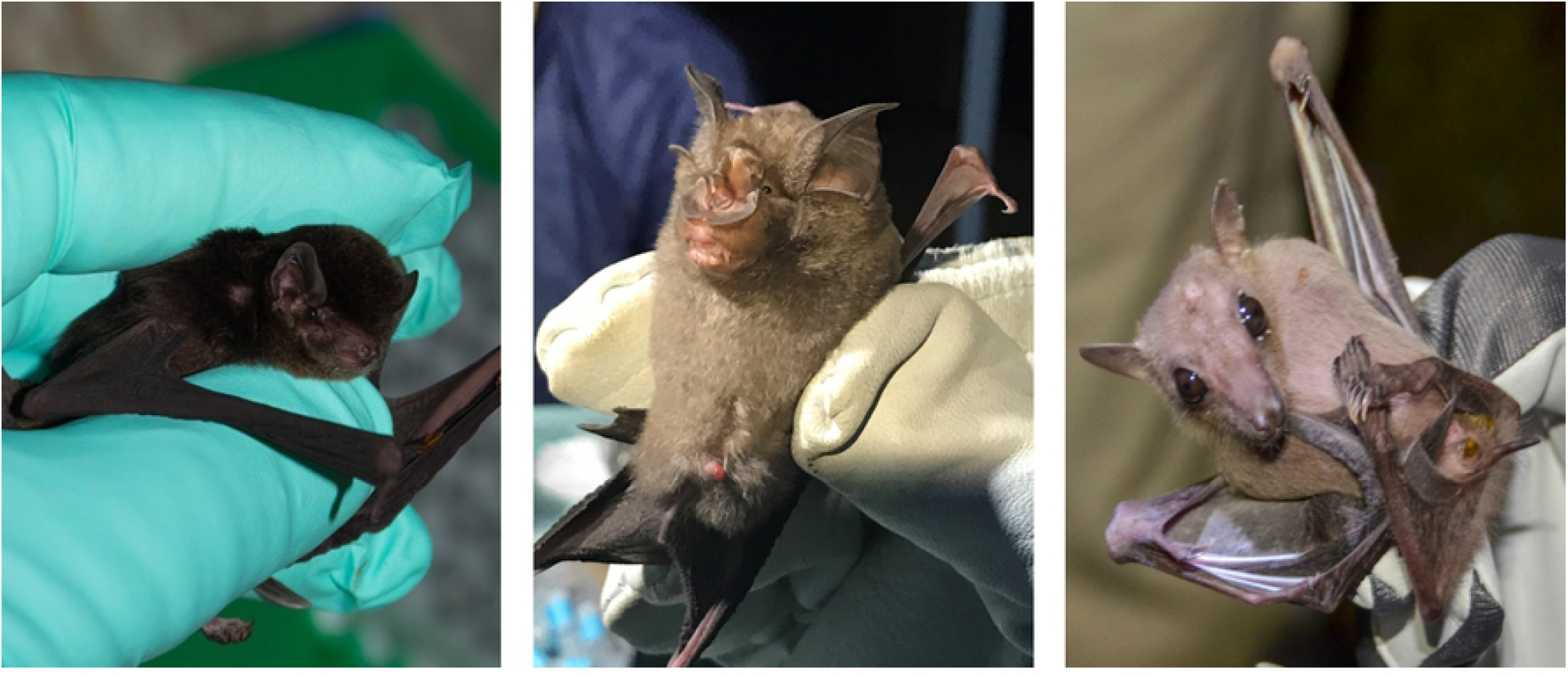
*Miniopterus fuliginosus, Rhinolophus rouxii* and *Rousettus leschenaultii* captured from Wavulgalge cave, Sri Lanka (capture credits: Andreas Nitsche).

**Fig 2.**
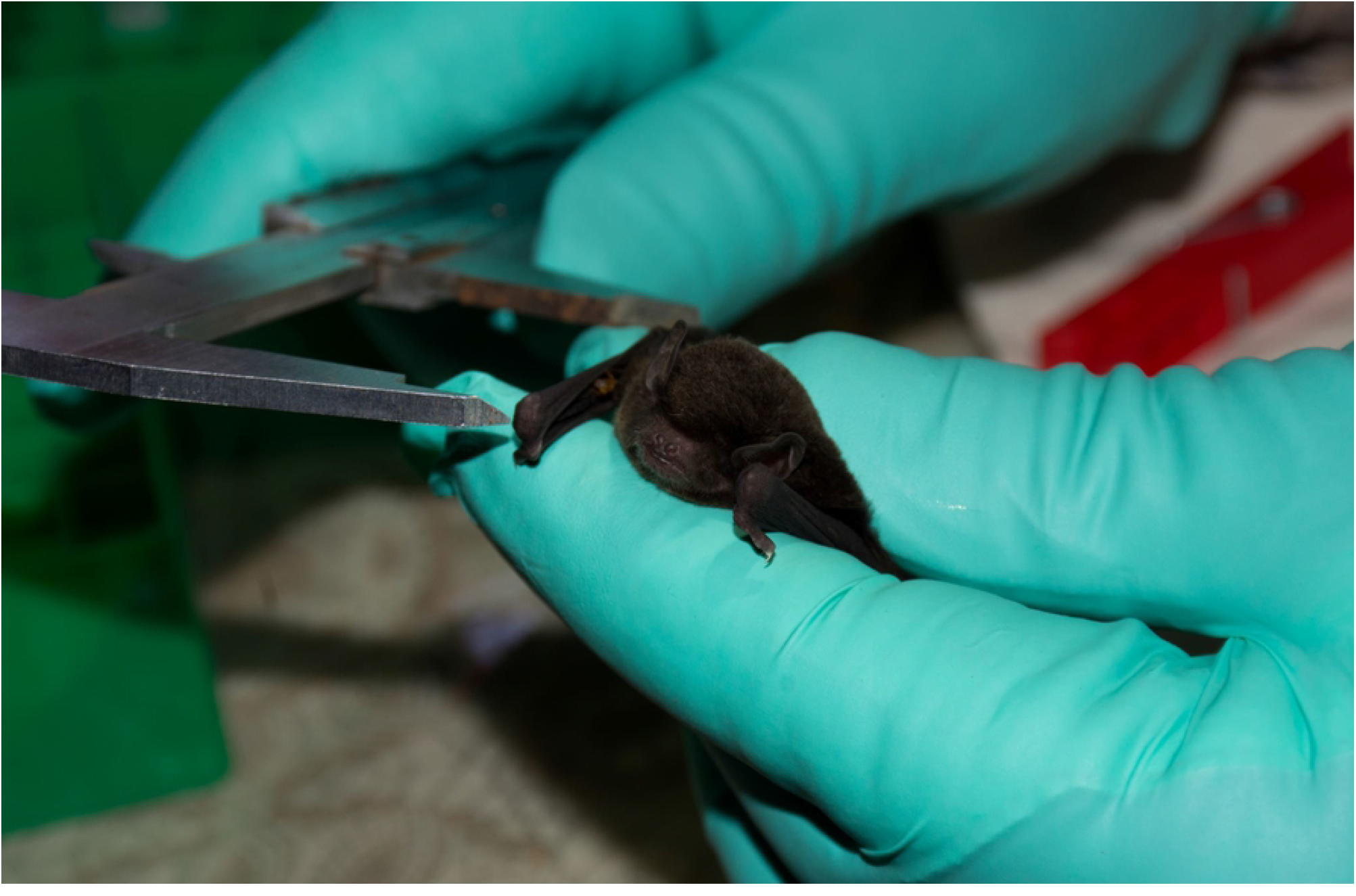
Collecting morphometric measurements. **Measuring forearm length of a bat using a vernier caliper (capture credits: Andreas Nitsche)**

An amount of 200 µl of Phosphate-buffered saline was added into each vial containing a dry oral swab and vortexed vigorously. QIAamp Viral RNA Mini kit (Qiagen, Hilden, Germany) was used to extract the nucleic acids from oral swab samples according to the manufacturer’s instructions. A 250 bp length of partial sequence of MT-CYB gene was amplified using FM up and FM do primers as described previously (9). Further, the full MT-CYB gene was amplified using RrFP and RrRP primers as previously described (10). PCR products were purified using MSB Spin PCRapace purification kit protocol (Invisorb®, Invitek GmbH, Berlin, Germany) following the manufacturer’s instructions. Purified products were sequenced on Applied Biosystems 3500 Dx Genetic Analyzer, using the corresponding forward and reverse primers for each strand. The BigDye™ Terminator v3.1 Cycle Sequencing Kit (Thermo Fisher Scientific, Waltham, MA, USA) was used to perform Sanger sequencing reactions.

The sequences obtained were analyzed using Geneious Prime® 2020.0.5 software. Sequences were trimmed and assembled using the De Novo Assembly function to obtain the consensus sequences. The consensus sequence of each bat was compared with the sequences in the database of NCBI using the Basic Local Alignment Search Tool (BLAST) to identify the respective species and calculate the statistical significance of matches. The results were compared with the initial morphological identification.

Further bioinformatic analysis was carried out using MEGA X 10.1 software. MT-CYB gene sequences (1,140 bp) of selected bat species were downloaded from the NCBI database and aligned with the consensus sequences obtained from this study using the MUSCLE alignment tool in the MEGA X application. The best substitutional model for the alignment was calculated using the, J model test (11). Phylogenetic reconstruction was carried out using the MrBayes Markov chain Monte Carlo (MCMC) method (Parameters were as follows: Substitution model, GTR, rate variation, equal; chain length 10 million; burn-in, 30%; sampling frequency, 200; *Felis catus* was used as an outgroup). Phylogenetic trees were visualized using FigTree software (http://tree.bio.ed.ac.uk/software/figtree/) and posterior probabilities were calculated for each node (12). A heat map was calculated based on the nucleotide alignment and percentage of identity.

## 3. Results

The obtained sequences are the first *mitochondrially encoded cytochrome b* full gene sequence data of Sri Lankan bats. Partial sequences (250 bp) of the MT-CYB gene were obtained for all bats sampled. Additionally, 74 full MT-CYB sequences of 1,140 bp length were obtained for *Rousettus leschenaultii, Rhinolophus rouxii, Hipposideros speoris, Hipposideros lankadiva and Miniopterus fuliginosus*. Table 1 provides all data collected for the 74 individual bats.

**Table 1.**
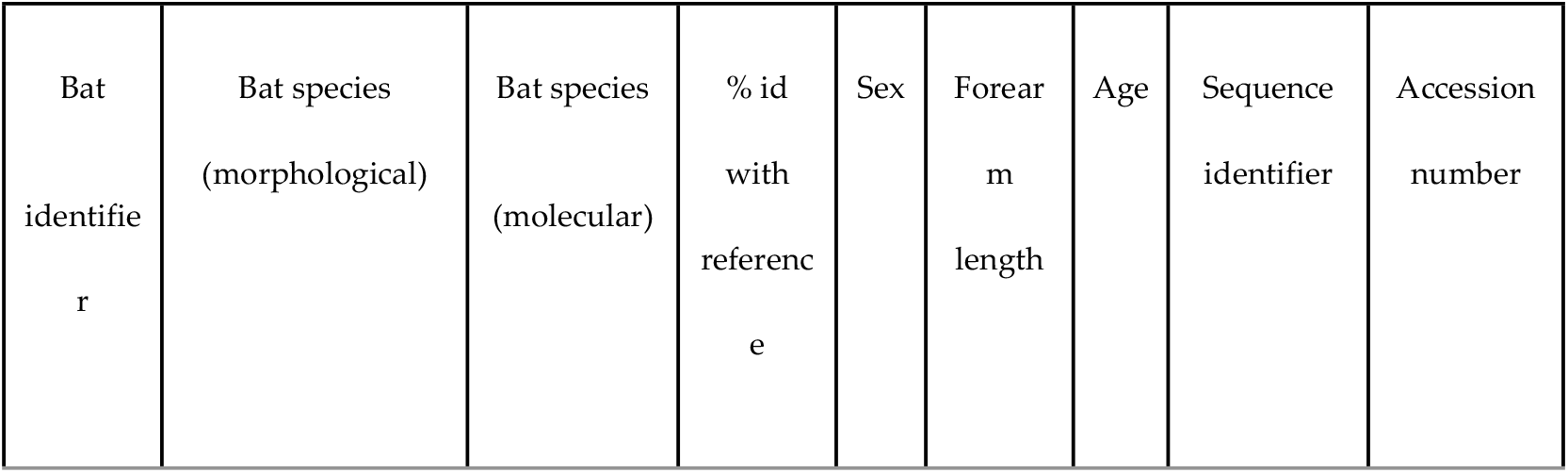
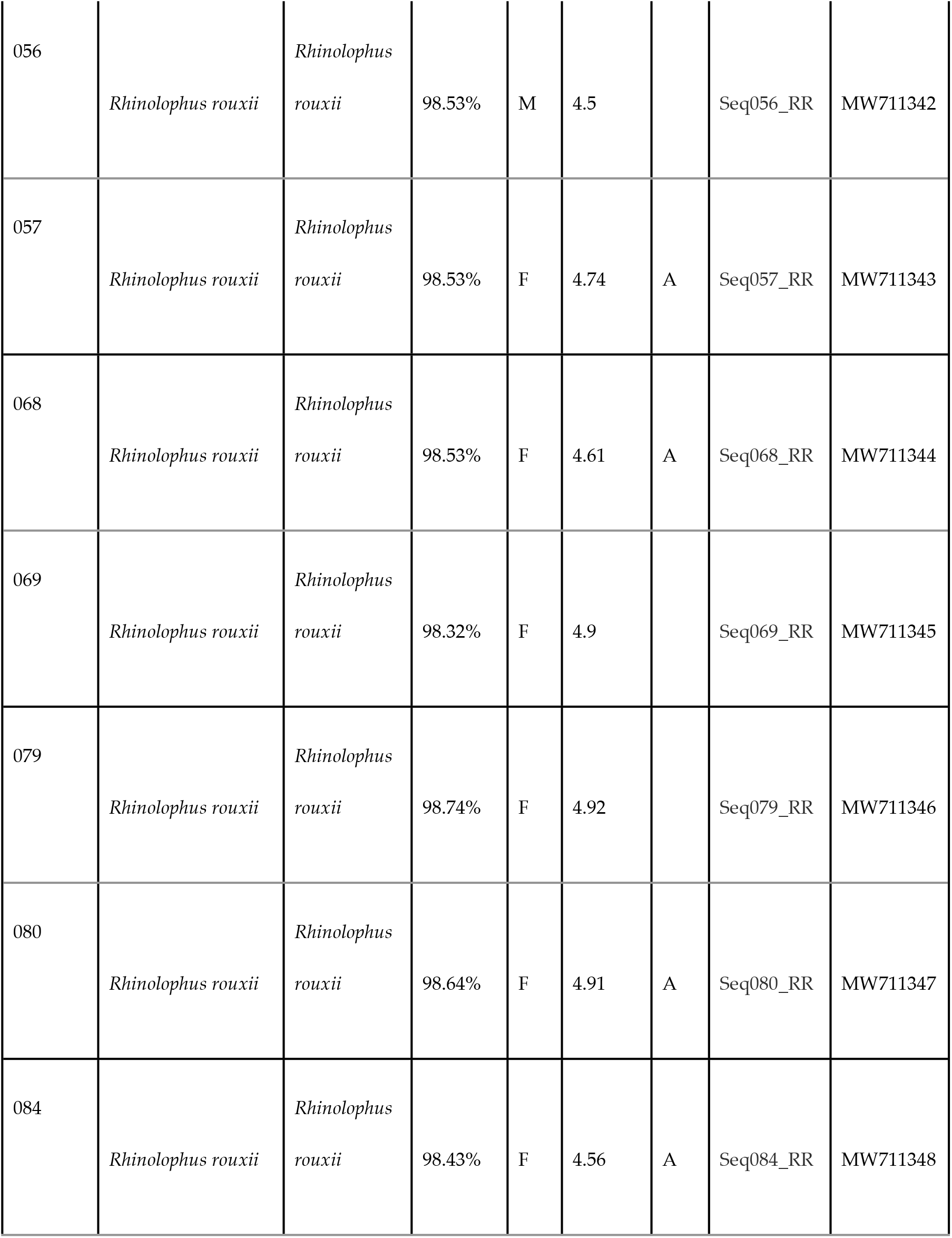
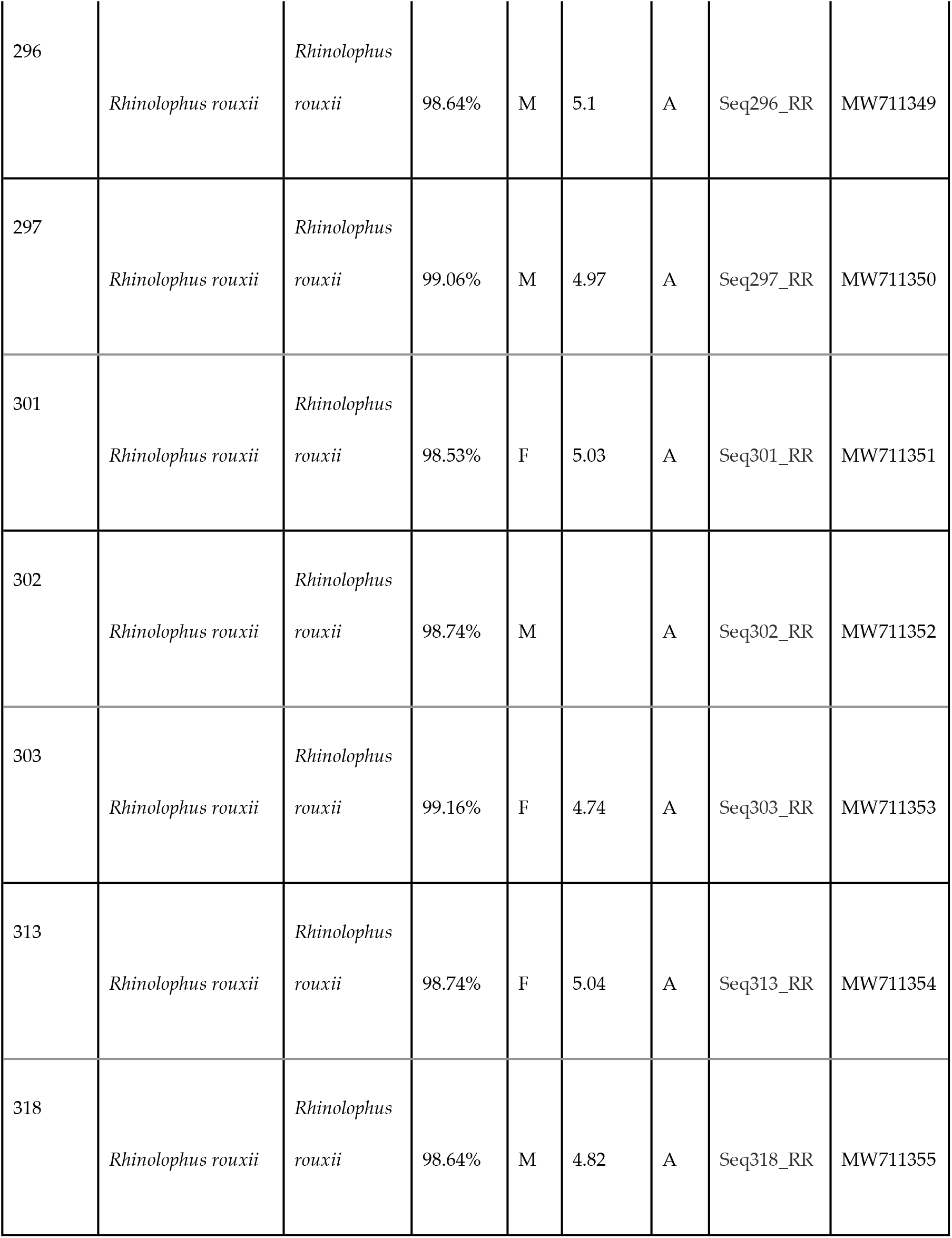
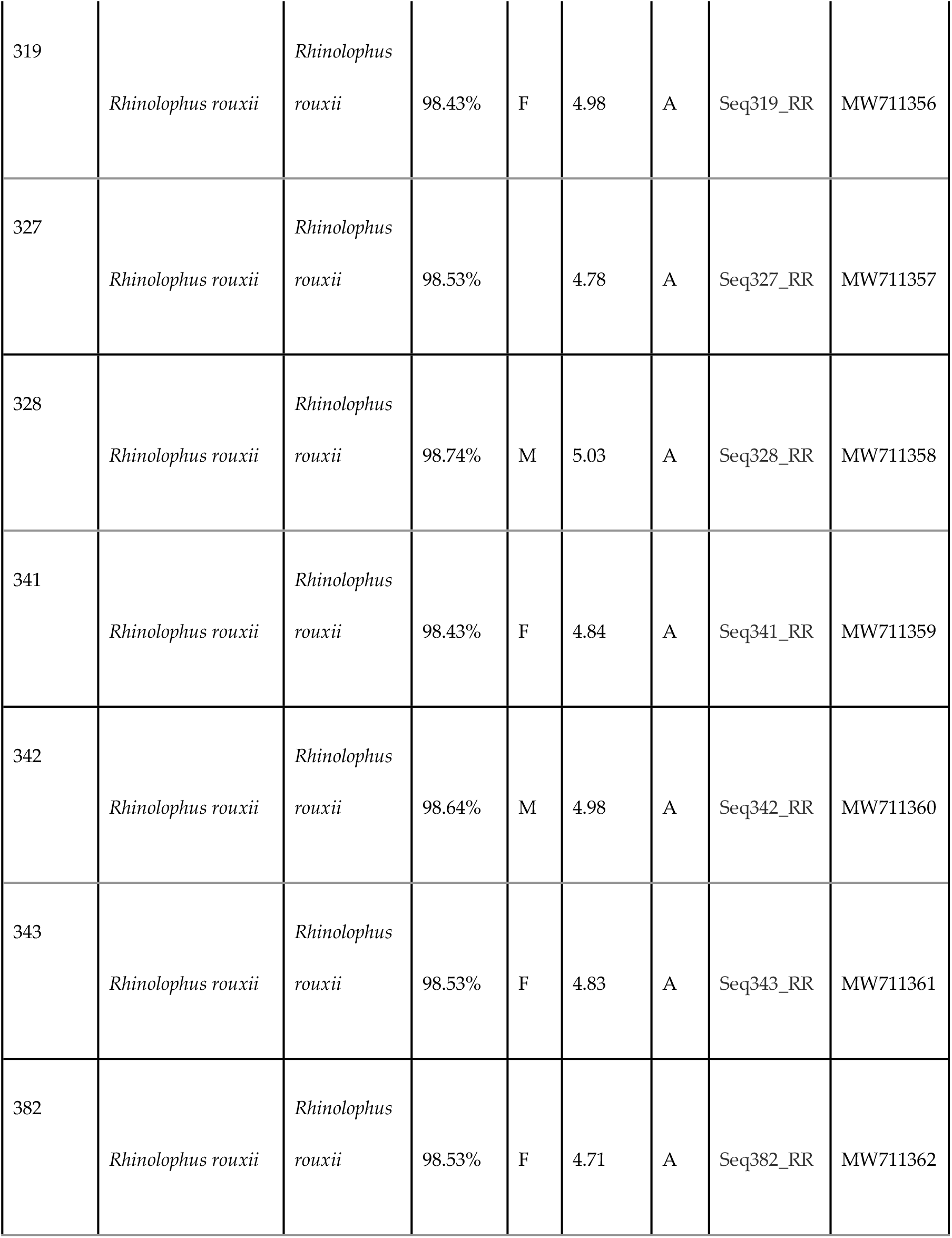
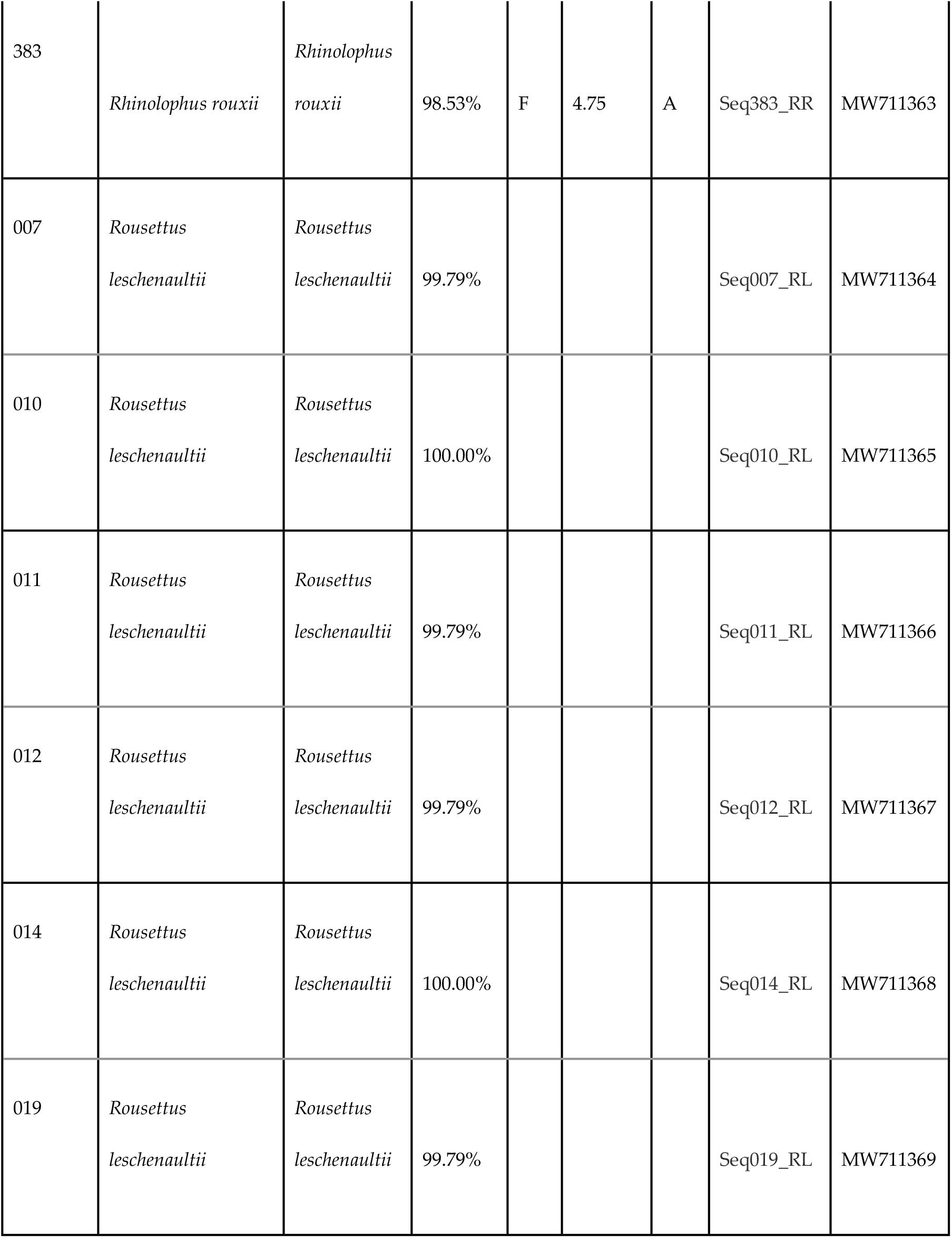
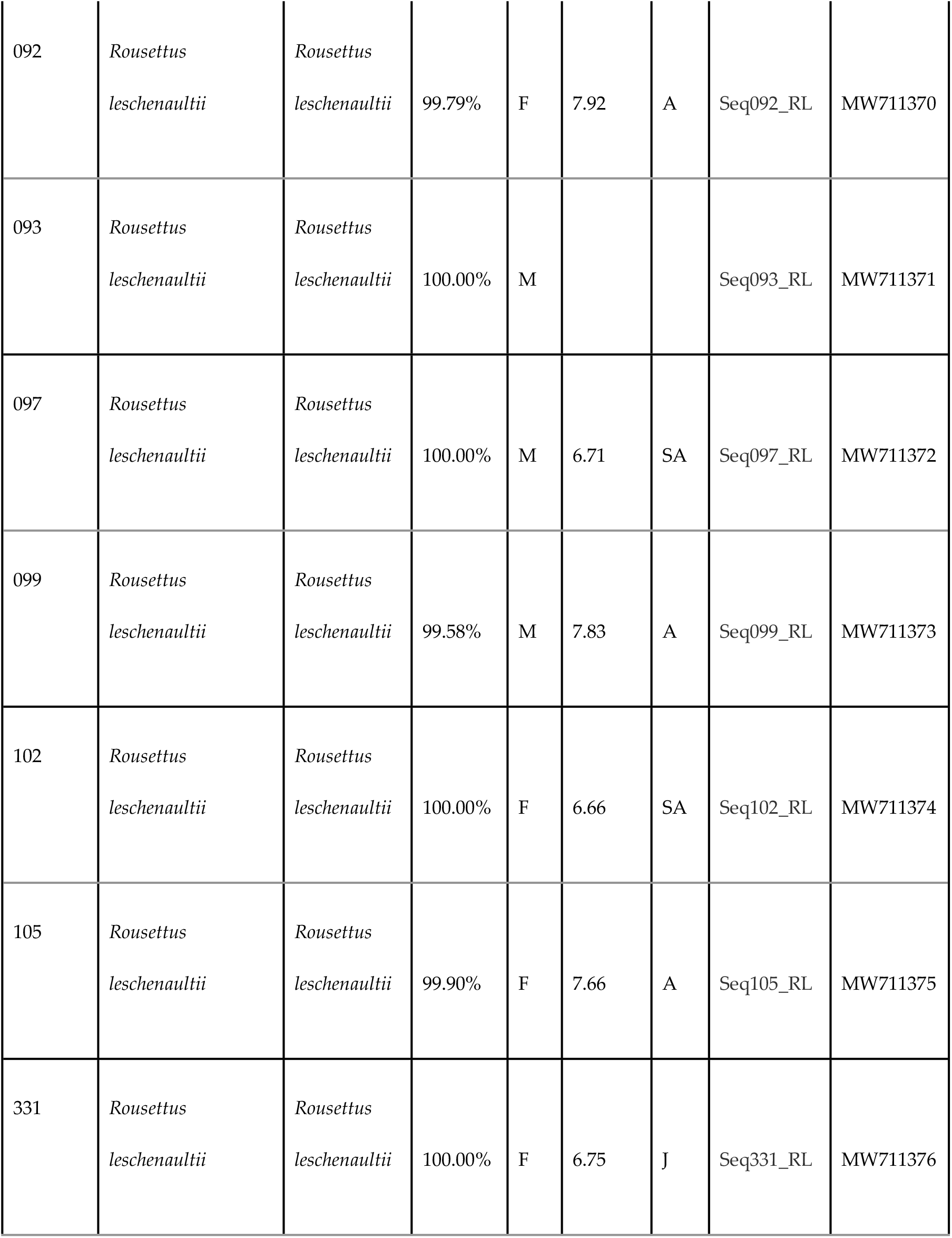
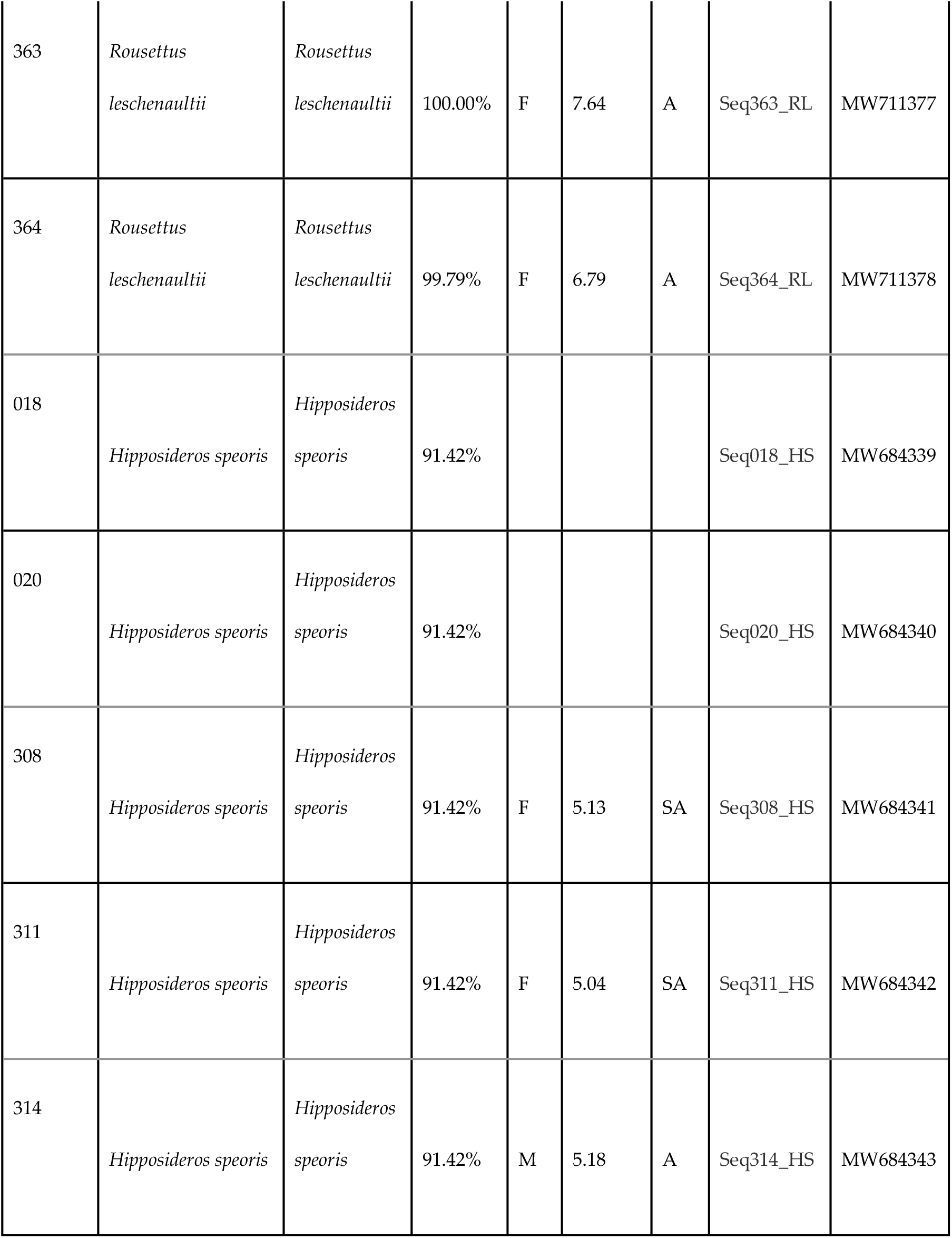
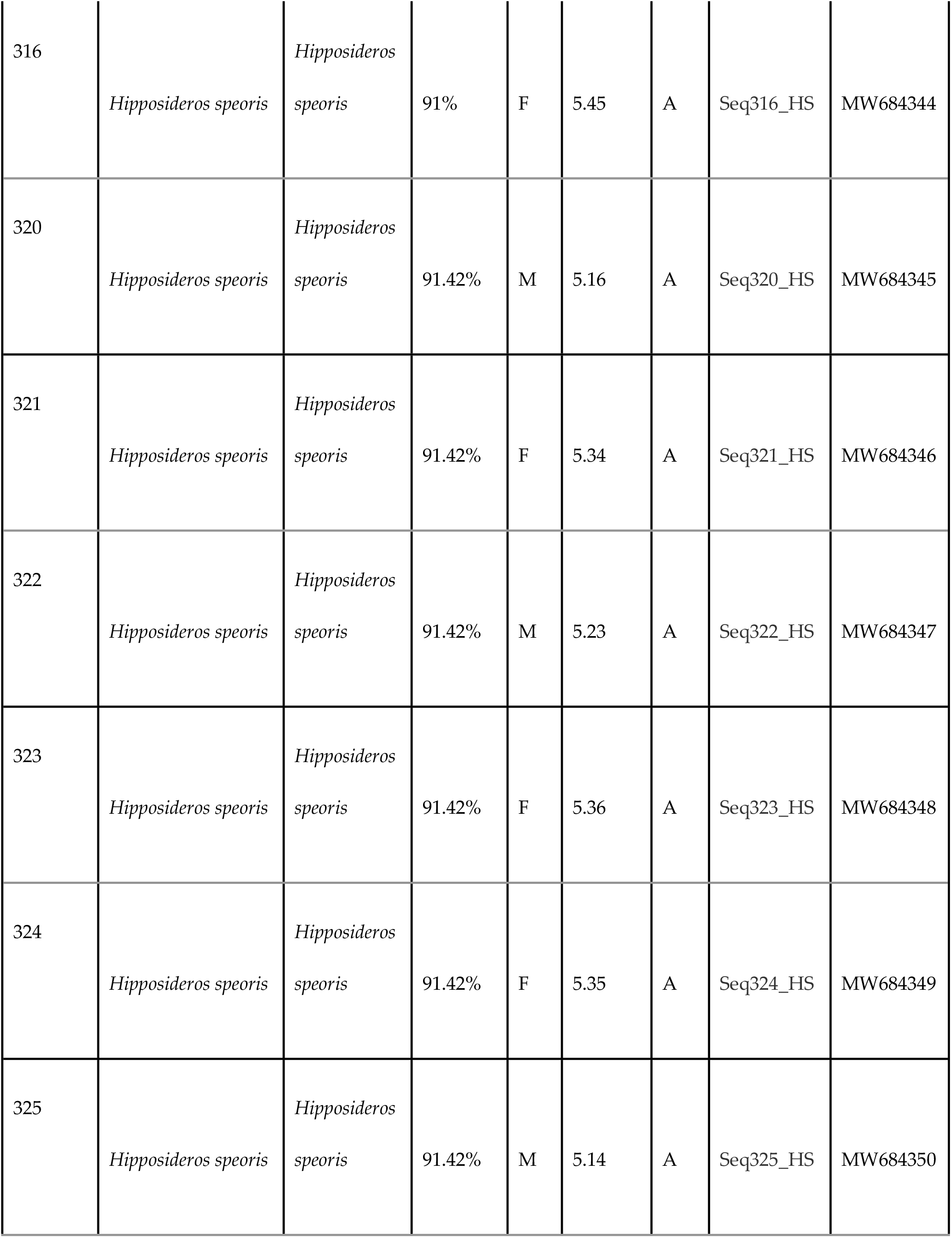
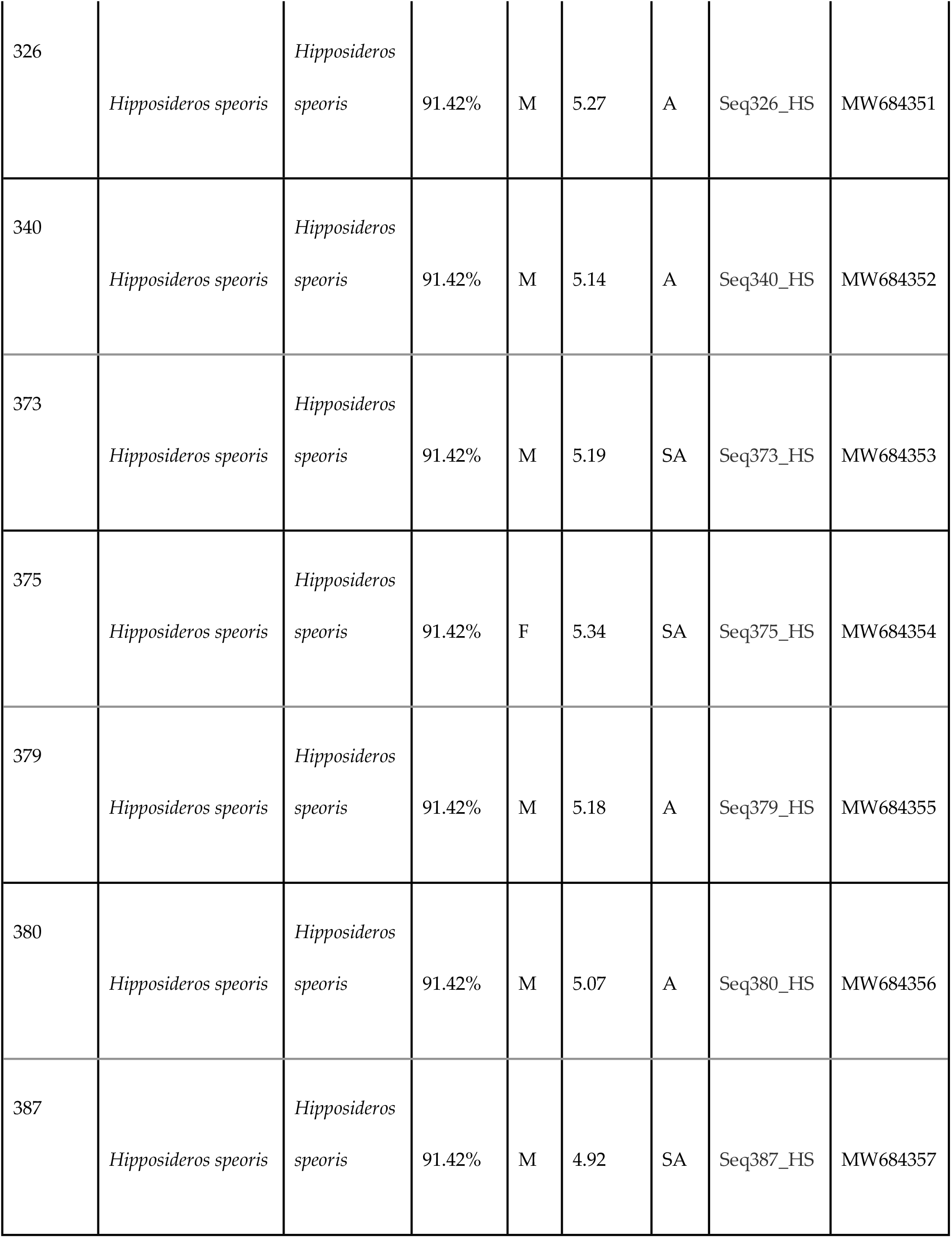
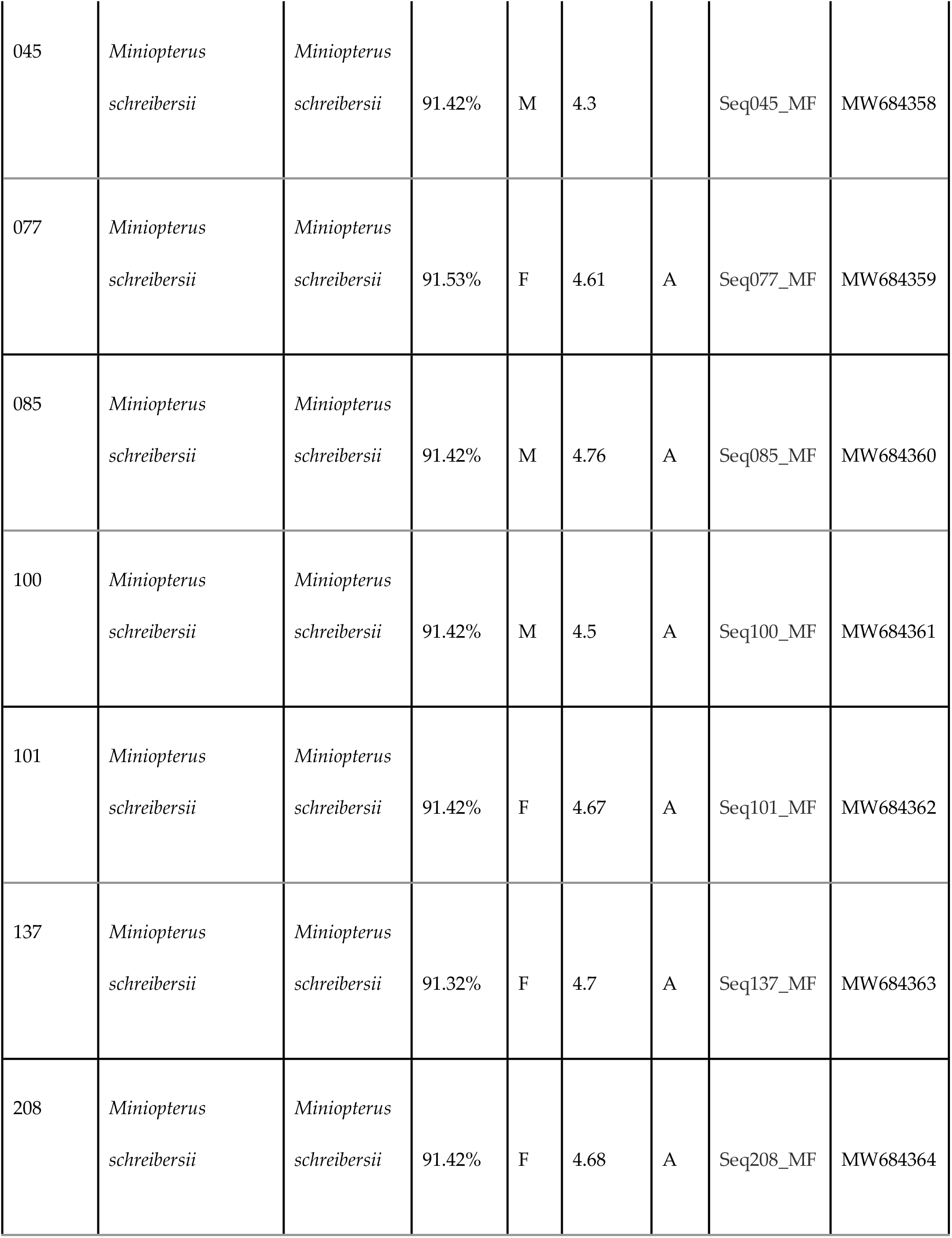
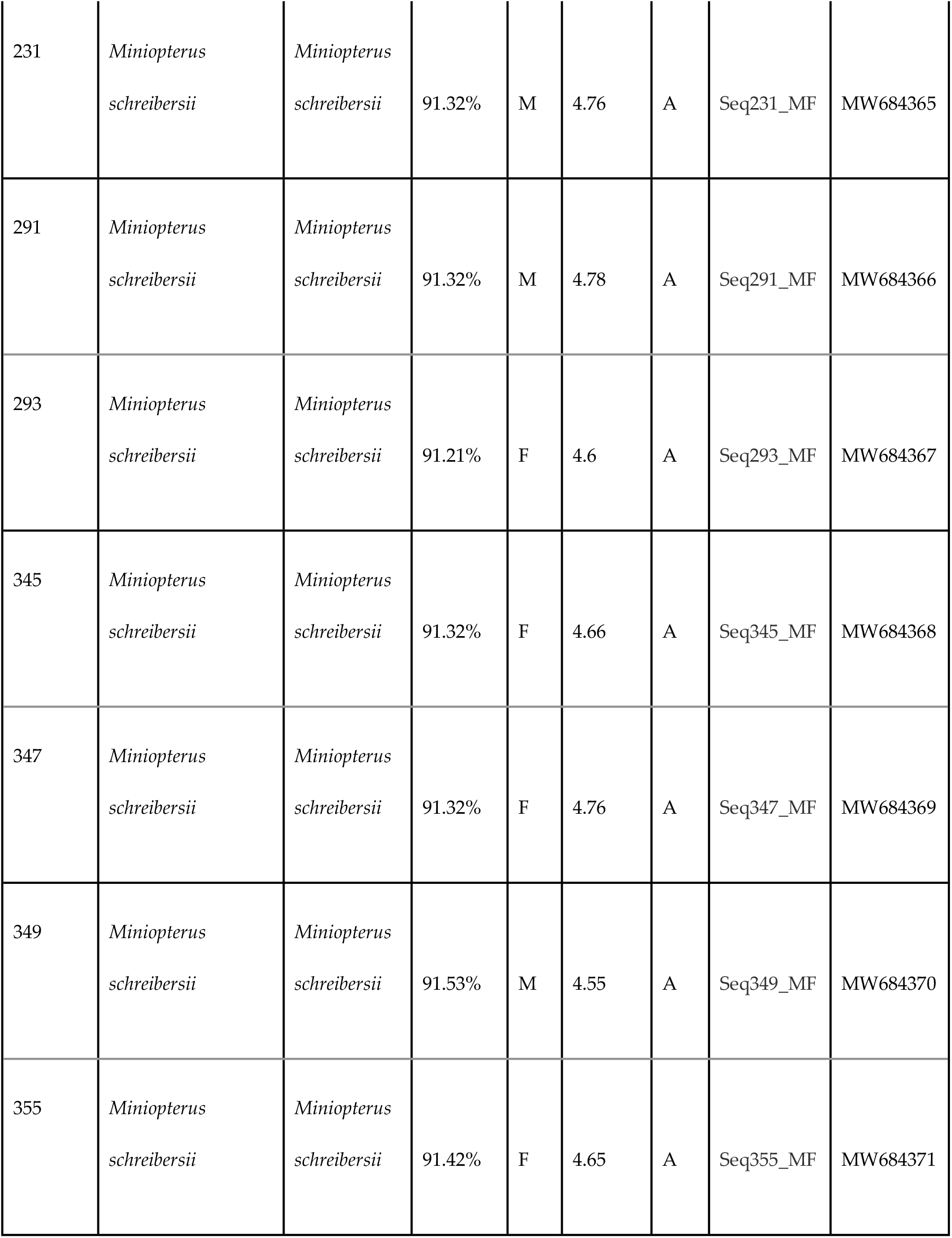
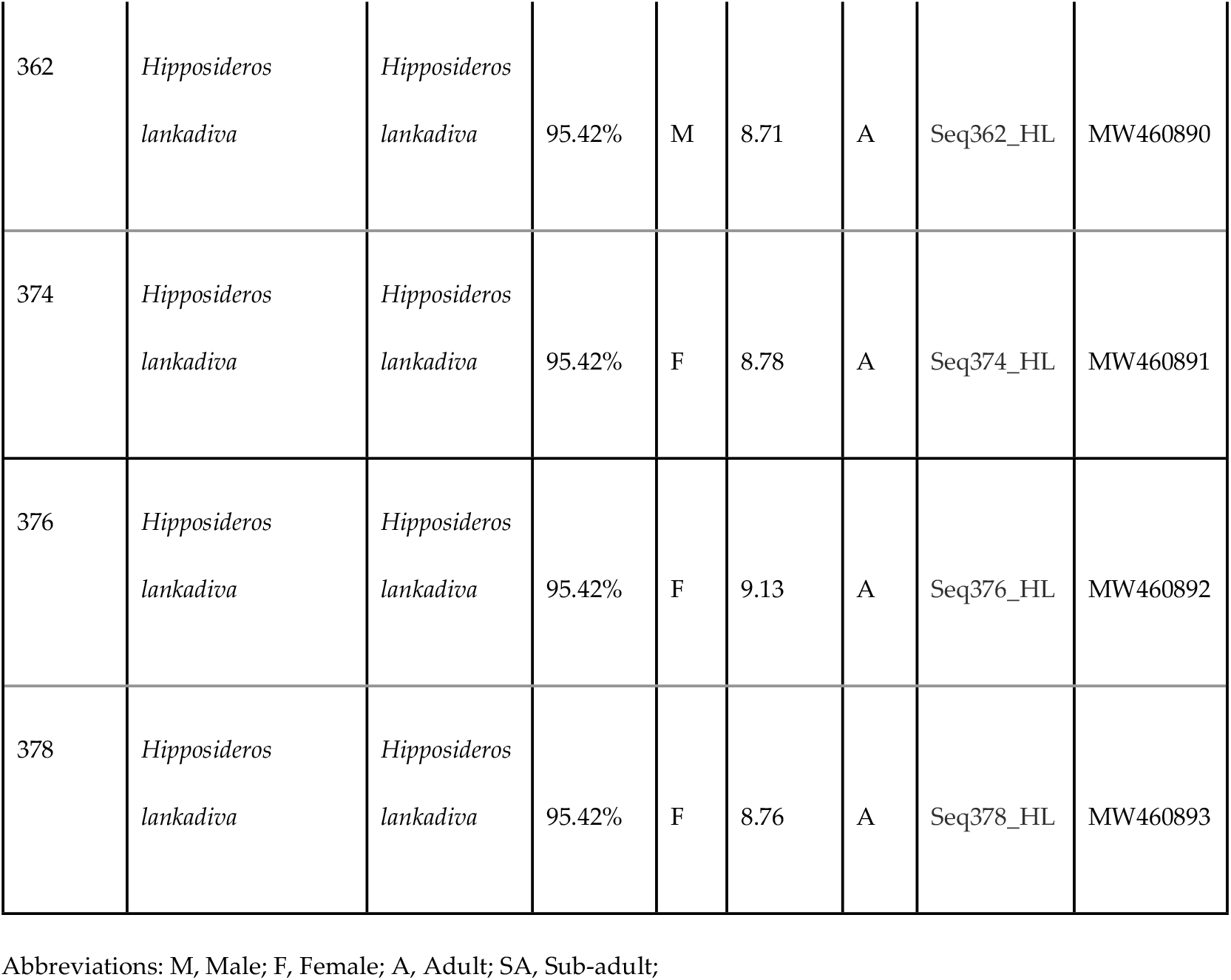
Overview on data collected for bats from Wavulgalge cave, Sri Lanka.

Phylogenetic trees are constructed using molecular data and used to make inferences to understand the evolutionary relationship among taxa. Phylogenetic reconstruction allocates the sequences of the five Sri Lankan bat species to the corresponding bat families within the species tree (Fig 3.).

**Fig 3.**
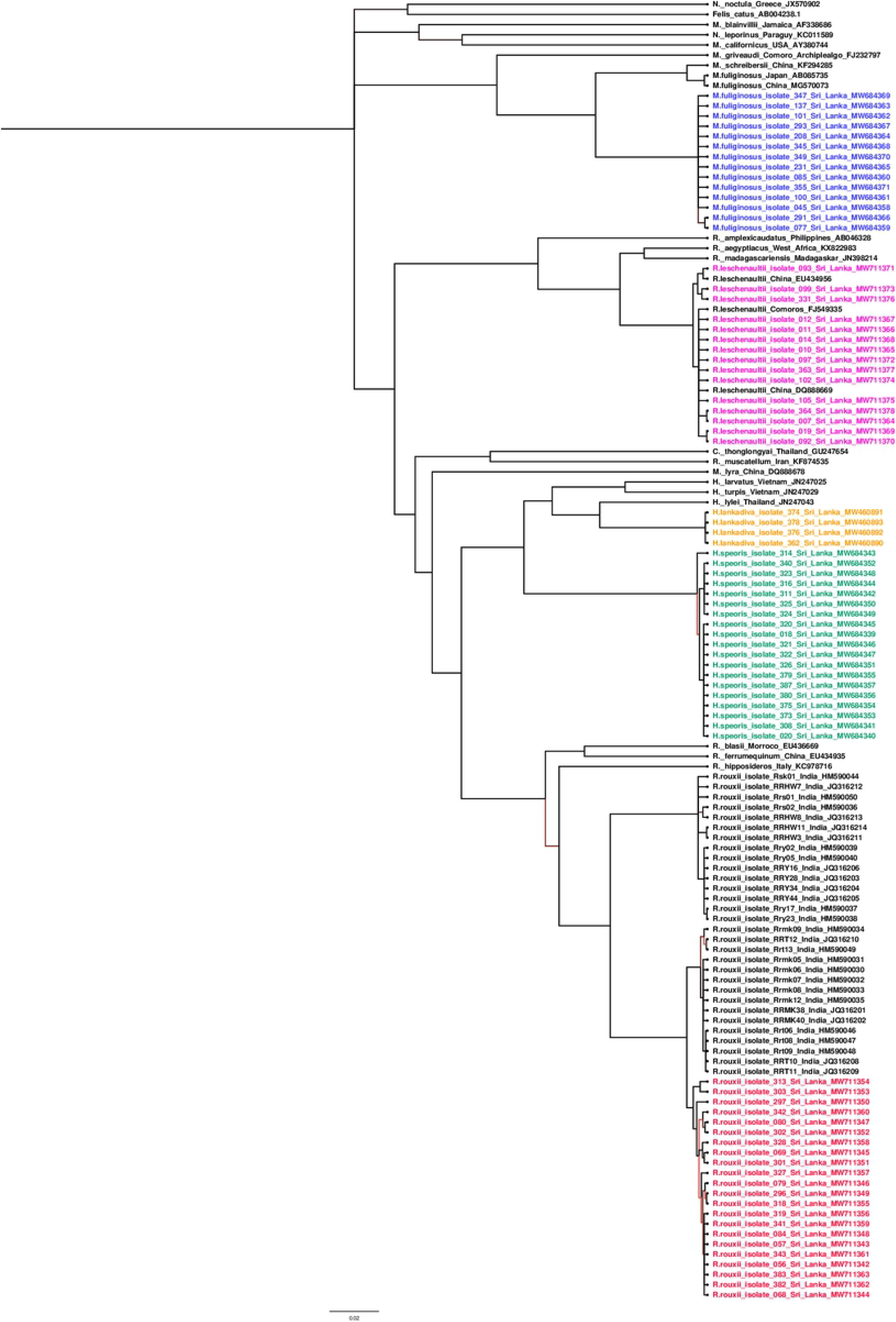
Phylogenetic reconstruction of bat species based on the full mitochondrially encoded cytochrome b gene (1,140 bp) in comparison to other species of bats. Reconstruction was calculated using MrBayes MCMC method (Parameters were as follows: Substitution model, GTR, rate variation, equal; chain length 10 million; burn-in, 30%; sampling frequency, 200; Felis catus was used as an outgroup). Reconstructed trees were visualized using FigTree and posterior probabilities were depicted for each node (http://tree.bio.ed.ac.uk/sofware/fgtree/).

*M. fuliginosus* and *H. lankadiva* show very low levels of intra-species variation. Sequences from *H. speoris* appear to divide into two distinct sister-clades. Sequences from *Rousettus leschenaultii* are also divided into two clades while showing a close relationship to sequences obtained from *R. leschenaultii* from China. Sequences obtained from *R. rouxii* are clustering as a distinct sister-clade to *R. rouxii* sequences obtained from India. According to the phylogenetic tree, Sri Lankan *R. rouxii* are more closely related to the *R. rouxii* species identified in Srirangapattana and Moodbidri regions than Yercaud and Sirumalai regions in India.

The relationships observed by phylogenetic reconstructions are further supported by the heat-map of distances, based on the percentage of genetic identity for all five bat species (Fig 4-7).

**Fig 4.**
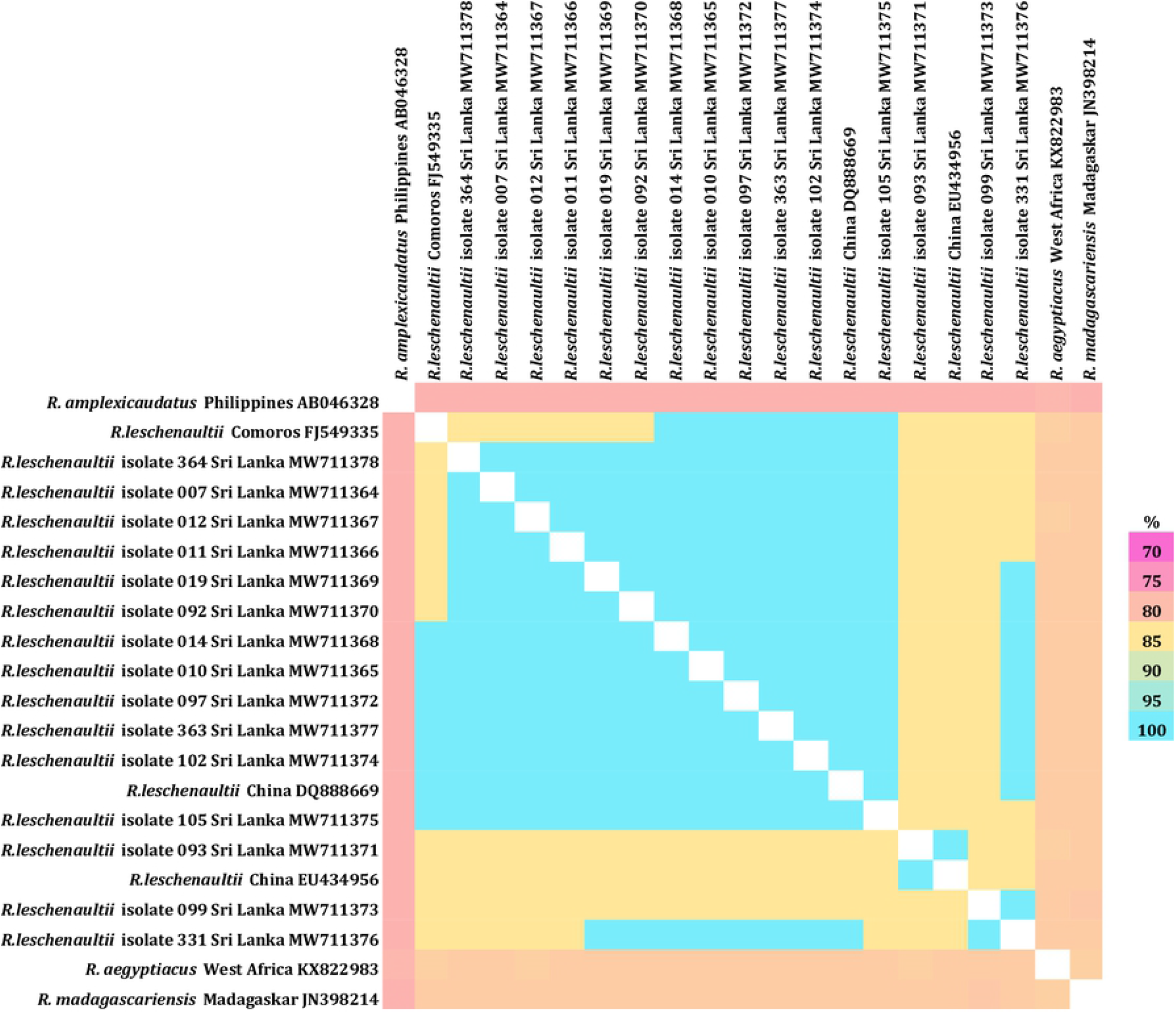
Heatmap based on the full MT-CYB gene (1,140 bp) of *Rousettus leschenaultii* in comparison to other species of bats. Percentage of identity is depicted by color ranging from 70 percent (red) to 100 percent (green).

**Fig 5.**
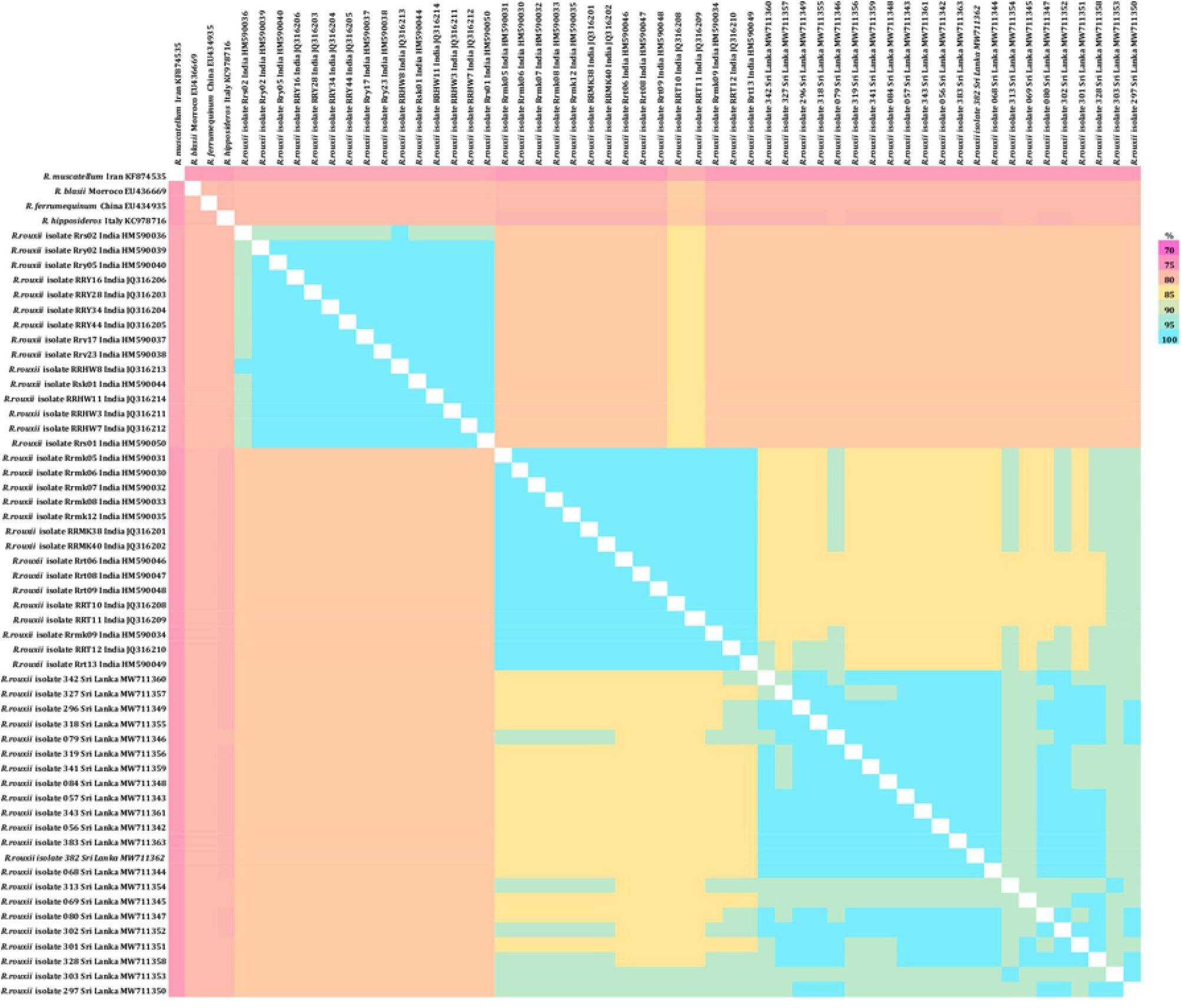
Heatmap based on the full MT-CYB gene (1,140 bp) of *Rhinolophus rouxii* in comparison to other species of bats. Percentage of identity is depicted by color ranging from 70 percent (red) to 100 percent (green).

**Fig 6.**
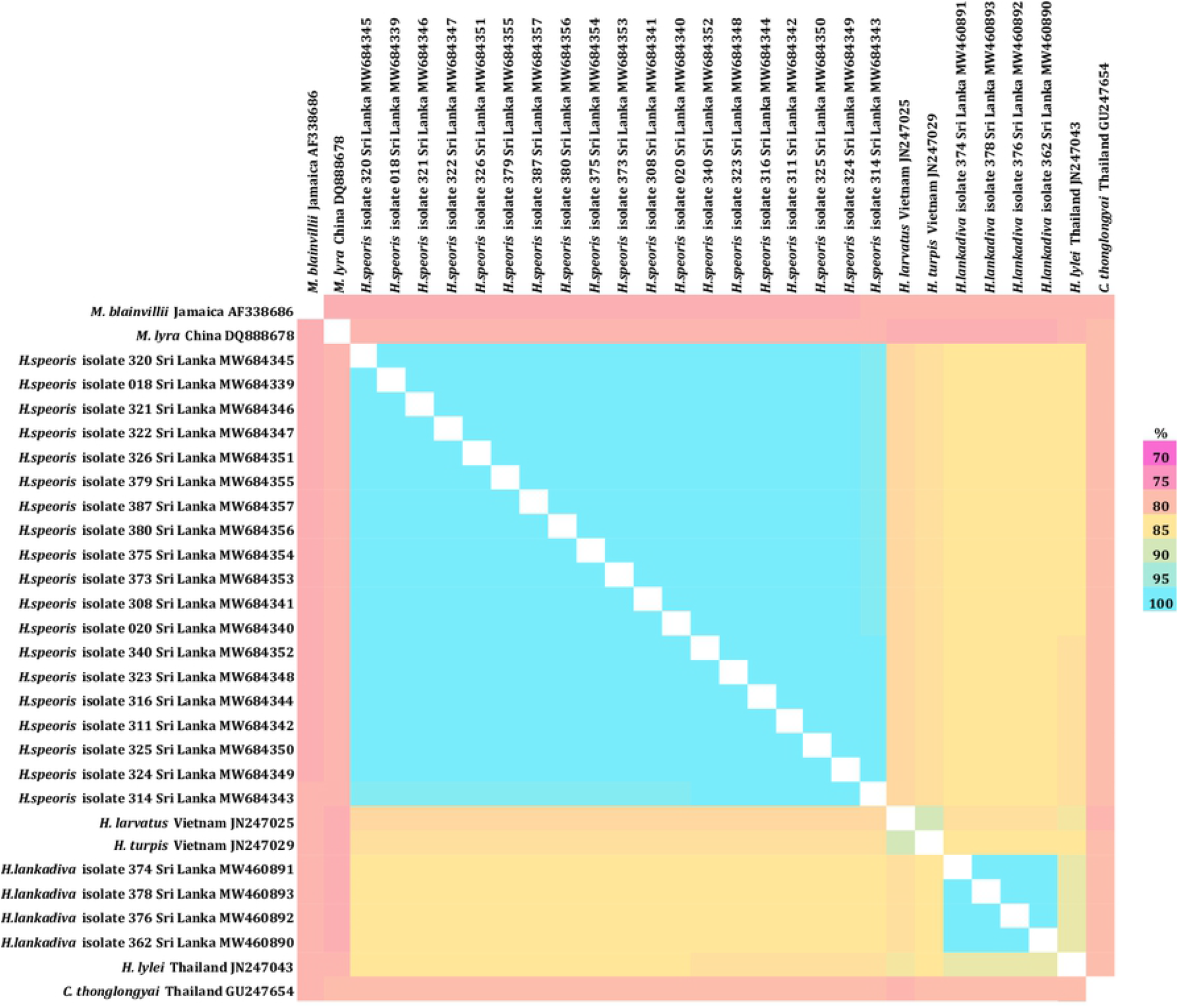
Heatmap based on the full MT-CYB gene (1,140 bp) of *Hipposideros speoris* in comparison to other species of bats. Percentage of identity is depicted by color ranging from 70 percent (red) to 100 percent (green).

**Fig 7.**
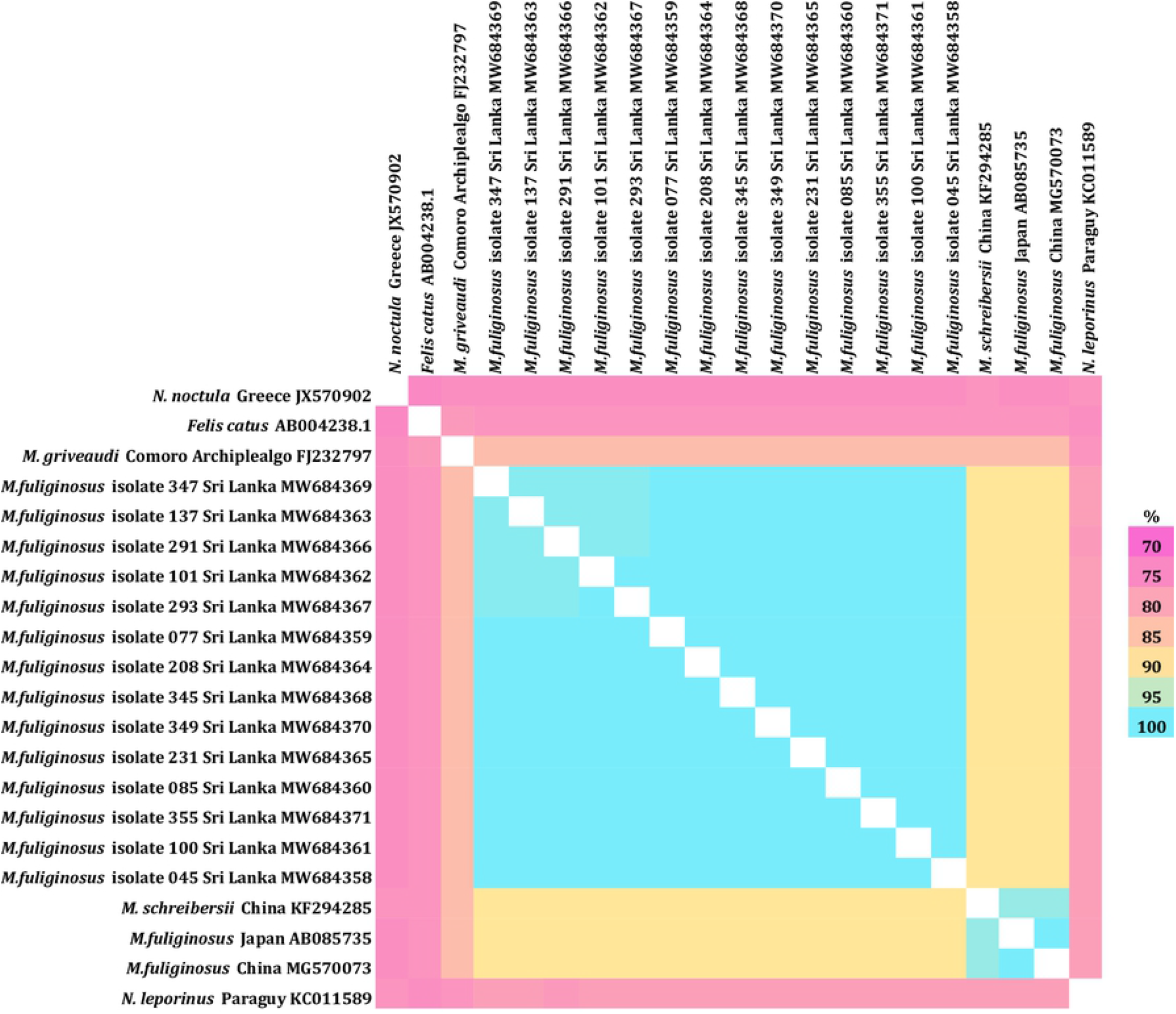
Heatmap based on the full MT-CYB gene (1,140 bp) of *Miniopterus fuliginosus* in comparison to other species of bats. Percentage of identity is depicted by color ranging from 70 percent (red) to 100 percent (green).

## 4. Discussion

Bat species in Sri Lanka were first described by Kelaart in 1852 (13). Later in 1935, Phillips carried out a more detailed study of mammalian species in Sri Lanka based on their morphological features (14). Species description of the Sri Lankan mammals by Phillips is still used as the reference guide for the identification of bat species in Sri Lanka (15). Many morphologically similar species have been recorded in neighbouring South Asian countries (10). With this study, we provide the first data for accurate molecular identification of five bat species inhabiting one of the largest sympatric bat colonies in Sri Lanka.

Further, we provide the first bat species tree for selected Sri Lankan bats based on MT-CYB genealogy. Our data analysis further supports the recent classification of bats into Yinpterochiroptera and Yangochiroptera suborders (4). Phylogenetic tree resulted in the monophyletic clustering of both *Hipposideridae, Rhinolophidae* and *Pteropodidae* bats and showed a clear divergence from the *Vespertilionidae* bats.

In contrast to other bat species inhabiting the cave, *M. fuliginosus* bats migrate to the Wavulgalge cave from nearby roosts during their maternity period. Thus, using the Wavulgalge cave as a maternity roost (16). However, the sequence analysis data show less variation between the individuals of *M. fuliginosus* bats, suggesting the robust similarity of the MT-CYB gene sequences of *M. fuliginosus* bats from different locations in Sri Lanka.

A total of four bats belonging to *H. lankadiva* were sampled during our field visits. Molecular data indicated that all four bats are highly similar to each other and more closely related to *H. lylei* (Thailand), *H. larvatus* (Vietnam) *and H. turpis (*Vietnam) bat species than *H. speoris* identified from Sri Lanka.

The phylogenetic tree revealed that the *R. rouxii* species from Wavulgalge cave is closely related to *R. rouxii* identified in the Karnataka district in India. There are two phonic types of *R. rouxii* recorded in India with echolocation call frequencies above 90 kHz and below 85 kHz3 (10,17). According to the phylogenetic data analysis from our study, it was revealed that Sri Lankan *R. rouxii* species is more closely related to the phonic type with echolocation call frequency below 85 kHz. Interestingly, the echolocation call frequencies of *R. rouxii* in Wavulgalge is below the echolocation calls of Rufous horseshoe bats recorded in India and fluctuate between 73.5kH −79 kHz (18).

*R. leschenaultii* sequences obtained in this study, appeared to be closely related to *R. leschenaultii* sequences from China. The phylogenetic reconstruction suggests that *R. leschenaultii* may have diverged from *R. aegyptiacus* and *R. madagascariensis* recorded in Africa. These three species are distinctly related to the *R. amplexicaudatus* recorded from the Philippines. This information is supported by a previous study carried out on *Rousettus spp*. phylogeny (19). In our phylogenetic reconstruction, *R. leschenaultii* appears in two distinct clades of the population. *R. leschenaultii* has the highest individual count for a single bat genus in Sri Lanka (2). It is also the most common species recorded in the Wavulgalge cave and occurs in all six bioclimatic zones of the island (15). Therefore, more extensive research should be conducted to determine the genetic diversity of *Rousettus* species in Sri Lanka.

It has been documented that, zoonotic pathogens which have co-evolved in bats have a higher risk of infecting other mammalian species, including humans (20). Bats are found to carry a multitude of different microorganisms that include viruses, bacteria, fungi and parasites (21). Furthermore, the global interest of bats as potential reservoir hosts of zoonotic pathogens highlighted recently as these flying mammals have been detected to be harbouring SARS CoV-2 (22). In addition, recent bat virome study revealed alpha-and betacoronaviruses in *Miniopterus fuliginosus* and *Rousettus leschenaultii* bats inhabiting the Wavulgalge cave, Sri Lanka (23). Prevalence of medically important pathogens such as SARS CoV-2 in bats may be linked with the bat species, their geographical location, roosting nature and other behavioural aspects (24,25).

Therefore, accurate species identification using molecular techniques will help to predict the pathogen-host shifts and interspecies pathogen transmission in bats in the Wavulgalge cave in Sri Lanka in future studies (23,26).

## 5. Conclusions

Our study addressed the research gap of the molecular taxonomy of Sri Lankan bats. Our results are consistent with the species identification by morphological identification of the bat species in the Wavulgalge cave. Accurate identification of bats plays a critical role in conservation. Therefore, our findings will also contribute to future conservation and systematic studies of bats in Sri Lanka. Further data collected from our study will provide the basis for a genetic database of Sri Lankan bats.

## Acknowledgments

We thank Angelina Targosz, Nicole Kromarek, Marica Grossegesse for technical and field assistance, RKI sequencing lab for providing us Sanger sequencing results. We convey our gratitude to the Department of Wildlife Conservation, Sri Lanka and the Institute of Biology, Sri Lanka for granting us the necessary permits.

## Financial Declaration

This study was partly supported by the Federal Ministry of Health, Germany (Bundesministerium für Gesundheit, BMG) under the IDEA (IDentification of Emerging Agents) project within the Global Health Protection Programme (GHPP).

## Data Availability Statement

The data presented in this study are openly available in GenBank (https://www.ncbi.nlm.nih.gov/genbank/).

## Author contribution statement

Conception and design of the project: TP, AN, WY, CK, SP, GP

Field work and sample collection: TP, TM, SS, BBZ, NK, WY

Performing experiments: TP, FS, TM

Analysis and interpretation of research data: TP, FS, JW, AN, WY, CK

Drafting of the manuscript: TP, CK

Critically revising significant parts of the work: AN, CK

Overall Supervising: SP, JW, SH, SF, GP AN, WY, CK

